# Application of eDNA metabarcoding for high-resolution reconstruction of the trophic web of an Arctic fjord

**DOI:** 10.64898/2026.01.09.698612

**Authors:** Chiara Piroli, Nadia Marinchel, Simone Galli, Tommaso Russo, Maurizio Azzaro, Francesco Filiciotto, Gabriele Di Marco, Alessia D’Agostino, Angelo Gismondi, Adriana Profeta

## Abstract

In the face of a rapidly changing Arctic, the ecosystem of Kongsfjorden was put under the spotlight to explore its community composition and structural dynamics. An eDNA metabarcoding approach was implemented to carry out a Food Web Analysis. eDNA samples were collected using *metaprobes,* innovative passive samplers, deployed under two different sampling configurations: in association with set fish traps in the coastal area and a towed sampling along a central transect, an offshore domain of the fjord. Amplification of the mithocondrial COI and ribosomal 18S genes was conducted in order to obtain a comprehensive view of metazoans and protists communities, respectively. The output taxa from the metabarcoding process constituted trophic webs’ nodes while producers-consumers and prey-predator’s relationships were identified through a literature review. Qualitative food networks were successfully obtained for each sampled site and for the two domains identified in the ecosystem, the coastal and offshore areas. Moreover, these networks were characterized by using four food web indicators: Species Richness (N), Number of links (L), Direct Connectance (C) and Generality (G). Differences in the apical part of the webs instantly emerged, as well as a clear separation between the coastal and offshore domain. Analyzing the values of the trophic indicators allowed for a deeper consideration regarding the nets’ structure and relative stability. Overall, eDNA proved sensitive and precise in capturing differences between the two domains and in providing insights into ecosystem structure. Moreover, eDNA-based Food Web Analysis could set the basis for long term monitoring studies in the same area, being cost-effective, rapid and easy to implement when compared to traditional methods.

## INTRODUCTION

Polar regions are widely regarded as ecologically fragile and particularly challenging to be studied due to a unique combination of biological sensitivity, rapid environmental change, and severe logistical constraints. Polar marine ecosystems are typically characterized by low functional redundancy, simplified food webs, and species that are highly specialized to extreme conditions of temperature, seasonality, ice cover, and resource availability. This high degree of specialization makes polar communities especially vulnerable to disturbance, as relatively small environmental changes or species losses can trigger disproportionate ecosystem-level responses (Clarke et al., 2005; Peck, 2018). Moreover, polar regions are experiencing climate change at rates exceeding the global average—most notably in the Arctic—resulting in profound alterations to sea-ice dynamics, ocean stratification, primary productivity, and species phenology, which in turn amplify the impacts of human pressures, such as fisheries, shipping, and tourism (IPCC, 2019; Meredith et al., 2019).

The rapid warming of the Arctic, with continued retreat of permafrost, snow and sea ice (Pachauri et al., 2014), is opening the region to more shipping, resource exploitation, port development, tourism, fisheries, and aquaculture, all of which have impacts on marine ecosystems (Ruiz & Hewitt, 2009). Fishing activities are a major source of disruption to marine food webs, both directly, by selectively removing components at specific trophic levels, and indirectly, by altering habitats and production cycles (Cicala et al., 2024). Additionally, Pecuchet et al. (2020) have stated that the introduction of boreal species in Arctic ecosystems (by poleward expansion due to climate warming) will affect ecosystem functioning and vulnerability, as these species will initiate feeding interactions with local residents, thus impacting food web properties. Altogether, it is expected a trend of homogenization of regional trophic networks, which will come at the expense of overall food web diversity being lost. It is therefore essential to monitor the status of marine communities in relation to climate change and anthropogenic activities.

Unfortunately, from a methodological and operational perspective, ecological research in polar marine environments is constrained by extreme weather conditions, limited accessibility, short sampling seasons, extended periods of darkness, and high logistical and financial costs. These factors have historically led to sparse spatial coverage and fragmented time series, reducing the ability to detect long-term trends, rare events, or cumulative impacts (Gutt et al., 2018). Consequently, polar regions serve as both early-warning systems for global environmental change and some of the most challenging settings for marine biodiversity monitoring, necessitating the development and application of innovative, integrative, and non-invasive approaches.

In polar marine systems, environmental DNA (eDNA) approaches—particularly metabarcoding of seawater (and, in some cases, sediments)—are increasingly used to generate non-invasive, high-throughput biodiversity baselines across taxa ranging from plankton and benthic invertebrates to fishes and megafauna, helping to fill major monitoring gaps in remote, logistically challenging environments. In Antarctica, nearshore eDNA metabarcoding has been shown to efficiently characterise metazoan communities and complements conventional surveys, while broader assessments have highlighted eDNA’s potential to expand the spatial and temporal coverage of Southern Ocean biodiversity monitoring. In the Arctic, eDNA metabarcoding has been proposed and tested as a surveillance tool for marine biodiversity in rapidly changing coastal systems, including under difficult seasonal conditions such as the polar night, thereby enabling more continuous ecological observation. Beyond baseline biodiversity mapping, eDNA is also emerging as a practical component of human-impact assessment and biosecurity, supporting early warning for non-indigenous species potentially associated with expanding shipping, tourism, and warming-driven range shifts; expert-driven frameworks explicitly highlight these applications for Antarctica, and recent field efforts demonstrate feasibility for detecting potential invaders in high-latitude routes and data-poor regions.

While its application to marine ecosystems is increasing, in the Arctic Ocean such type of investigation remains limited (Havermans et al. 2022). In the Arctic, eDNA studies took advantage of different genetic markers targeting various taxonomic groups from seawater (Merten et al., 2023) to sediment (van den Heuvel-Greve et al., 2021) and sea-ice samples (Leasi et al., 2021), with collection methods tailored to sample type (Ruppert et al., 2019).

In addition, eDNA-based approaches can be used to support Food Web Analysis (FWA; D’Alessandro and Mariani, 2021) as it allows the evaluation of the community structure and overall relationships between biodiversity and ecosystem functioning (Cicala et al., 2024). Trophic webs describe the connections among species according to their ecological interactions, such as predator–prey and producer–consumer (Smith and Smith, 2012; Le Guen et al., 2025). Moreover, assessing trophic relationships can help predict the relative stability of the ecosystem, when this is confronted with changes that can affect its structure (Renaud et al., 2011). Food web indicators have been developed to describe different ecological properties that can be interpreted as a response to an environmental or anthropogenic change (Le Guen et al., 2025).

Traditionally, methods of trophic web reconstruction include stomach content and stable isotope analyses. Even though these methods constitute robust models, accounting for carbon flows and biomass estimates (D’Alessandro and Mariani, 2021; Le Guen et al., 2025), they require specific taxonomic expertise, in addition to being time consuming and expensive. Moreover, they require a large sampling effort and are lethal to the sampled organisms (Le Guen et al., 2025).

In this context, the development of DNA metabarcoding-based approaches has opened up new scenarios for the rapid and large-scale collection of information on the presence and distribution of marine species (Cicala et al., 2024).

Within this framework, custom and self-produced samplers, called *metaprobes* (Maiello et al., 2022), have proved to be efficient passive eDNA collectors in the marine environment, consistently capturing overall community composition, as reported by studies from the Mediterranean Sea (Maiello et al., 2022; Maiello et al., 2024) and tropical reef ecosystems (Maiello et al., 2025).

Recent studies have demonstrated that combining eDNA data with literature-derived predator–prey interactions enable the reconstruction of realistic marine trophic networks (D’Alessandro and Mariani, 2021; Le Guen et al., 2025). Despite the absence of biomass data, eDNA-derived networks can still provide valuable insights into community structure and ecosystem stability, which could help environmental scientists and practitioners to monitor a larger portion of our seas and flag possible ongoing anthropogenic disturbance (D’Alessandro and Mariani, 2021; Cicala et al., 2024). This makes eDNA metabarcoding a powerful and practical approach for food web reconstruction and ecosystem monitoring, especially in remote and extreme environments such as the Arctic. Its cost-effectiveness, non-invasiveness, and easily replicable natures are advantages when monitoring a rapidly warming region.

Kongsfjorden is a glacial fjord located in the European Arctic, on the west Spitsbergen coast in Svalbard. As an open fjord with no sill at its entrance, it is strongly influenced by the inflow of warm Atlantic Water from the West Spitsbergen Current, mixed with Arctic water on the shelf (Walczowski et al., 2012). Since 1994, oceanographic monitoring has shown that Atlantic Water enters Kongsfjorden annually, typically during summer (Hop and Wiencke, 2019). However, since 2006 this phenomenon has also been observed in winter (Cottier et al., 2007). This shift, driven by regional warming, reduced sea ice cover, and altered wind regimes, has intensified Atlantic inflow, a process referred to as Atlantification (Cottier et al., 2007; Hop & Wiencke, 2019). Owing to its position at the interface of Arctic and Atlantic oceanic regimes, the Kongsfjorden is considered an early warning indicator of future changes, which can then be extrapolated to a pan-Arctic scale (Hop et al., 2002). Kongsfjorden has a long tradition of research and monitoring, which has established it as a reference site for marine ecological studies and long-term observations, and it now serves as a key location for investigating the impacts of environmental change on Arctic coastal ecosystems (Hop and Wiencke, 2019).

The present study is framed within the broader context of the VECNA project, funded by CNR-IRBIM and aimed at detecting Non-Indigenous Species (NIS) presence in Kongsfjorden. Additionally, eDNA analyses were incorporated into the project, with samples collected using *metaprobes* to investigate the community composition of the ecosystem. Indeed, this study marks the first deployment of *metaprobes* in an Arctic environment, providing a pilot application of their use, as well as a test of their efficiency in a polar environment.

This research aims to evaluate the potential of eDNA metabarcoding, and, specifically, the effectiveness of *metaprobes*, as a tool for characterizing the ecosystem of Kongsfjorden. A multimarker metabarcoding approach was implemented, selecting the mitochondrial cytochrome oxidase subunit I (COI) and the 18S SSU ribosomal V8-V9 region, targeting metazoans and protists respectively, to get a comprehensive view of the ecosystem. Specifically, the main objective was the reconstruction and description of trophic networks, to improve understanding of community structure and ecological interactions. The nodes of the food webs were defined based on taxa identified through eDNA metabarcoding, and a literature review was conducted to investigate the trophic interactions among them. Overall, this approach highlights the potential of eDNA metabarcoding in the field of trophic ecology and ecosystem monitoring and its effectiveness in detecting differences inside marine communities’ structure, in a rapidly changing Arctic ecosystem.

## MATERIALS AND METHODS

### Sampling

The sampling activity was carried out in Ny-Ålesund, Kongsfjorden, Svalbard, in July 2023. Four nearshore sites, hereafter referred to as “coastal”, were selected: near Ny-Ålesund harbour (NNH); outside the west pier of the harbour (EP – “External harbour pier”); Kapp Guissez (NWK - north-western side of the entrance of the Kongsfjorden); Kvadehuken (SWK – south-western side of the entrance of the Kongsfjorden) (Figure 1 and Table 1). Additionally, four sites (labeled as TB stations) were identified along the central part of the fjord, and the corresponding samples were collected, hereafter referred to as “offshore” (Figure 1). The environment in the central part of the fjord differs from the pier area in front of the settlement of Ny-Ålesund, mainly in depth ranges: from 220 to 260 m approximately in the fjord to shallower waters around the shoreline.

**Figure 1.**
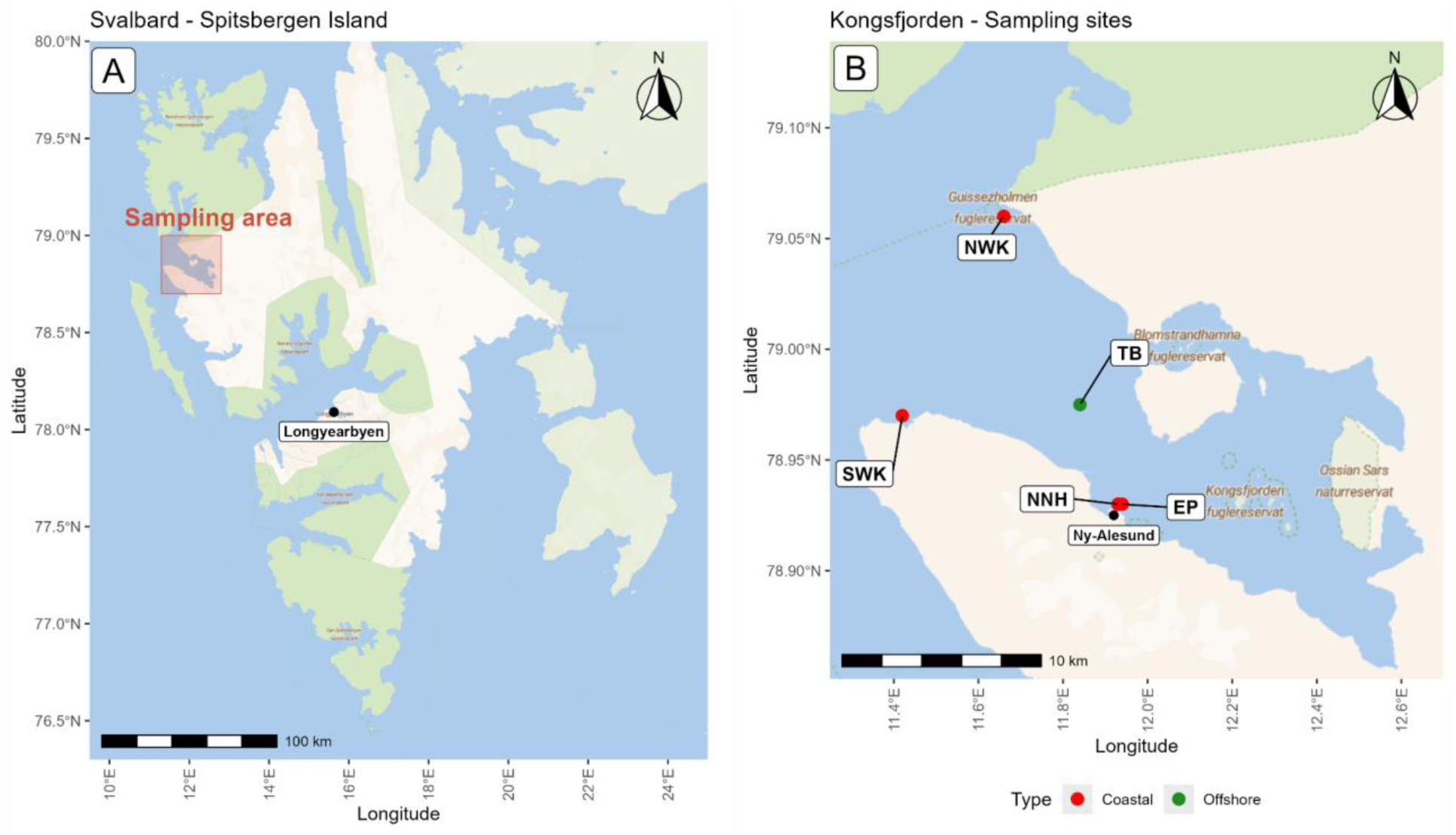
(A) The island of Spitsbergen, with the study area highlighted in red. (B) Sampling sites in the Kongsfjorden area: red dots represent coastal stations while the green dot is the mean point of the towed transect made by boat.

**Table 1.**
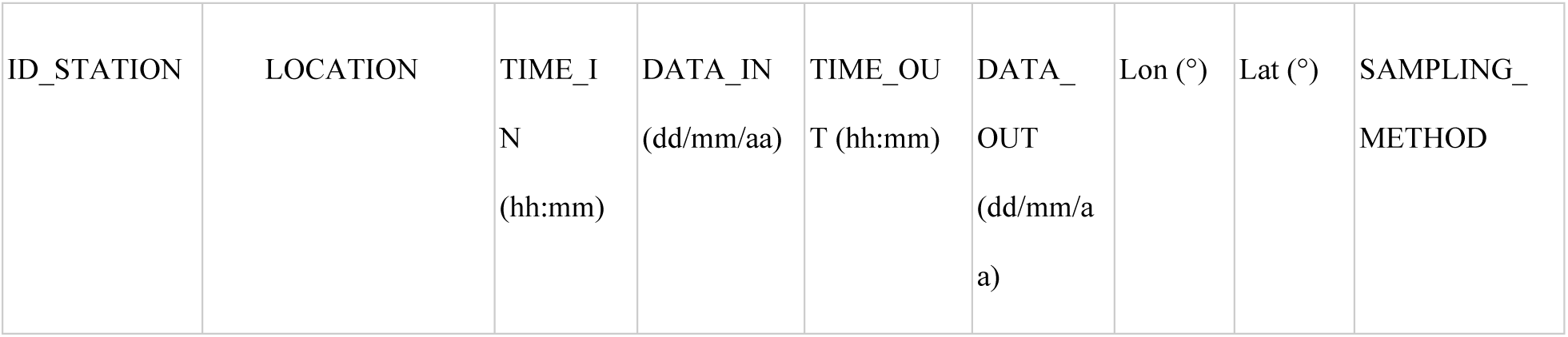
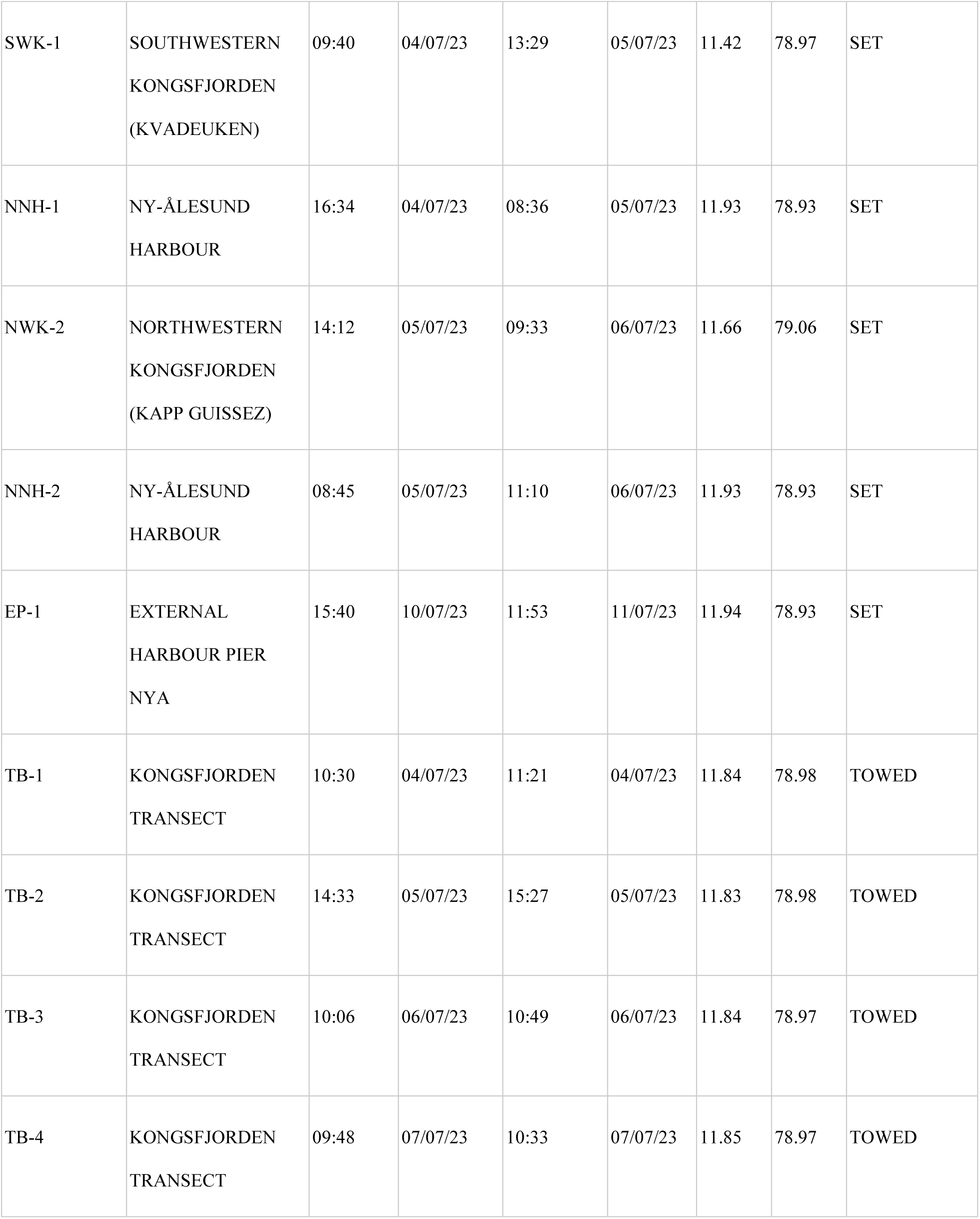
Overview of sampling stations in Kongsfjorden, including location, sampling times, coordinates, and sampling methods.

eDNA samples were collected using *metaprobes* (Maiello et al., 2022), i.e., 3D-printed bespoke hollow spheres which contain sterile gauze rolls tightly fixed by plastic cables, designed to passively capture DNA. Detailed instructions to produce and set up *metaprobes* are available at https://github.com/GiuliaMaiello/Metaprobe-2.1.

Samples were collected by means of two different strategies. Fish traps with a total length of 100 cm were used in the coastal locations. They were weighted with local rocks and placed at a depth varying between 2 to 6 m and linked to buoys for their identification and recovery. To attract fish, baits of commercial fish were inserted inside the traps. *Metaprobes* were then fixed inside fish traps, which were deployed for approximately 24 hours. This sampling strategy will be referred to as “set”, since the traps were static during their time in the water. For offshore samples, *metaprobes* were towed at the sea surface along the central part of the fjord at a speed of approximately 3.6 knots. This sampling method was noted as “towed”.

The rolls of gauze inside the *metaprobe* were then collected and placed in 50 ml sterile tubes containing 99% ethanol. Fish traps, tools to collect gauzes and *metaprobes* were cleaned after use and rinsed in a bleach solution to remove any traces of DNA. Samples were then stored at −20 °C in the laboratory before DNA extraction.

In total, nine samples were collected: five coastal and four offshore samples (Table 1).

### eDNA processing

#### DNA extraction

Laboratory processing of the samples was conducted at the University of Rome Tor Vergata. DNA was extracted from the gauzes using a custom protocol. Initially, a small section (∼2×2 cm) from the outer layer of the roll was cut into small pieces using sterile scissors. The pieces of gauze were placed in a 1.5 mL tube to allow the ethanol to evaporate. Each sample was then incubated at 56 °C for 24 h in 1 mL EDTA (0.5 M pH 8) and 0.25 mg/mL Proteinase K. After lysis, samples were centrifuged, supernatant was collected, and DNA was extracted with Roche Assembly Tubes silica columns (Dabney and Meyer, 2019). For each sample, two technical replicas were made; and a negative control of extraction for each batch (group of samples handled together in the extraction laboratory) was processed along with the samples.

#### DNA amplification and sequencing

DNA extracts were quantified using a Thermo Scientific NanoDrop Lite spectrophotometer and used as a template for polymerase chain reaction (PCR) amplifications targeting two genes: cytochrome oxidase I (COI) and 18S ribosomal RNA gene targeting the V8–V9 region (18S) (Table 2). Illumina adapters were added at 5’ end of the primers (forward adapter: TCGTCGGCAGCGTCAGATGTGTATAAGAGACAG; reverse adapter: GTCTCGTGGGCTCGGAGATGTGTATAAGAGACAG). PCRs were performed using a Bio-Rad iQ5 thermocycler in a 50 μL reaction volume containing the following components: 10 μL of template DNA, 2.5 U of JumpStart REDAccuTaq LA DNA Polymerase (Sigma-Aldrich, high-fidelity), 20 mM of each primer (Bio-Fab research srl), 0.2 mM of each dNTP (Sigma-Aldrich), 1X LA DNA Polymerase buffer (Sigma-Aldrich), 3 mM MgCl₂, and 5% DMSO. Amplifications were carried out under the following conditions: an initial denaturation at 95 °C for 5 minutes; followed by 35 cycles of denaturation at 95 °C for 20 seconds, primer annealing at 49.2 °C for 20 seconds, and extension at 72 °C for 35 seconds. A final extension step was performed at 72 °C for 10 minutes. Each set of samples included negative PCR controls.

**Table 2.**
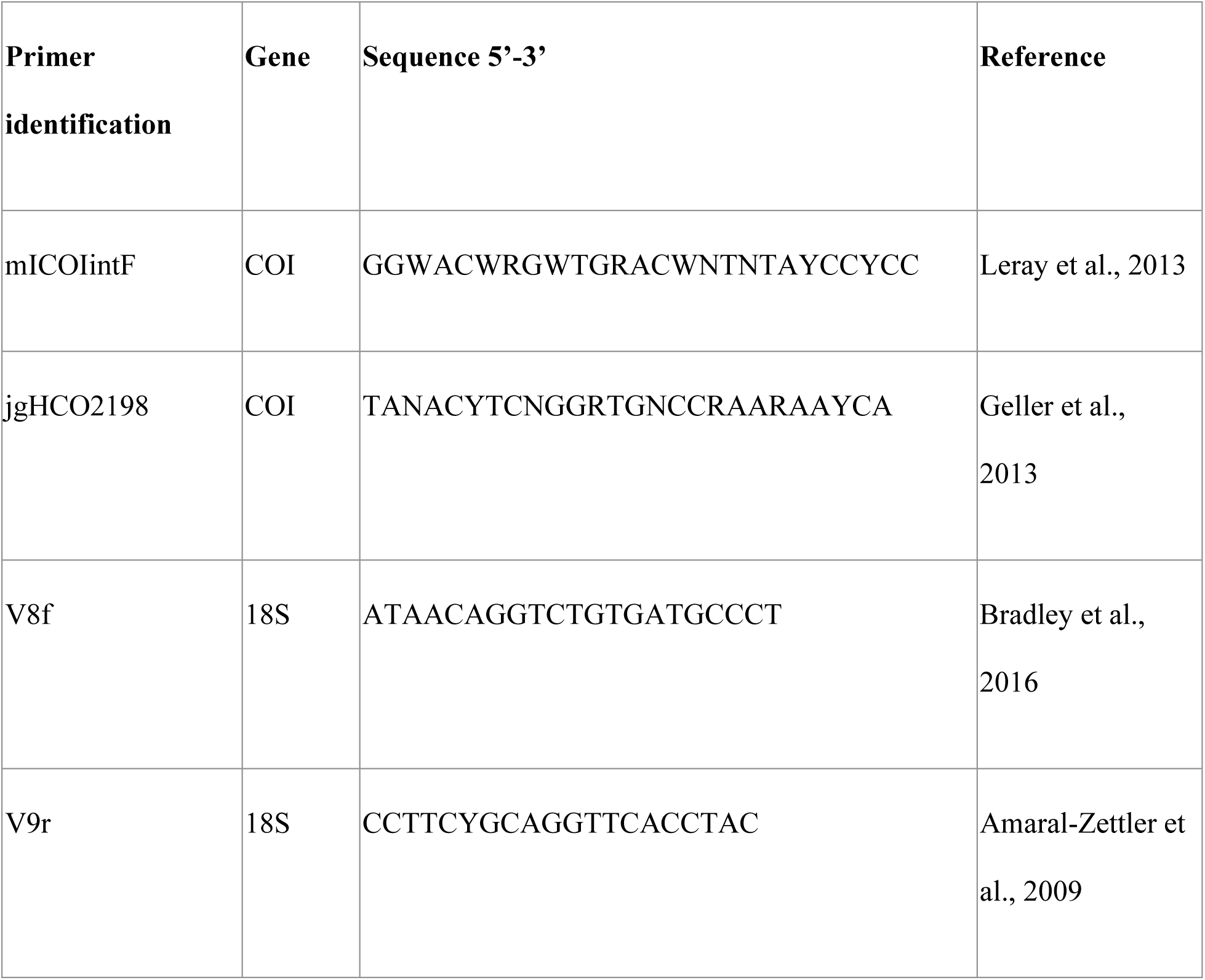
Primers used for eDNA metabarcoding, with target gene, nucleotide sequence and corresponding reference.

PCR products were verified in quality and quantified by a Thermo Scientific NanoDrop Lite spectrophotometer and sent to Bio-Fab Research s.r.l. (Rome, Italy) for Next Generation Sequencing. Library was prepared as follow: 1x PCRBIO HiFi Buffer (PCR BIOSYSTEMS, USA) containing 1 mM dNTPs and 3 mM MgCl2, 0.5 units of PCRBIO HiFi Polymerase (PCR BIOSYSTEMS, USA). Amplicons were then purified using MagSi-NGSPREP Plus magnetic beads (Euroclone, Milan, IT) and subsequently barcoded with Illumina® DNA/RNA UD Indexes, Tagmentation kit. Libraries were checked and quantified using the Tape Station 4200 (Agilent Technologies) and Qubit Fluorometer (Invitrogen Co., Carlsbad, CA) instruments and then pooled together such that each index-tagged sample was present in equimolar amounts. Sequencing was carried out on an Illumina NextSeq 2000 platform using a 2×300 bp paired-end run.

### Bioinformatics

Bioinformatic analyses were conducted using the OBITOOLS software (Boyer et al. 2016) for both COI and 18S raw reads. Firstly, forward and reverse primers were removed using CUTADAPT. Read quality assessment was performed with FASTQC, and low-quality bases were trimmed using OBICUT before downstream processing. Paired-end reads with a quality score exceeding 40 were merged using ILLUMINAPAIREDEND, while dereplication was performed with OBIUNIQ, which clustered identical sequences. Sequence filtering was performed using OBIGREP to remove ambiguous bases and reads falling outside the expected length range (290 - 340 bp for COI and sequences less than 350 bp for 18S, as the expected barcode length was of 313 bp for COI and 330 bp for 18S (Leray et al., 2013; Bradley et al., 2016)). Chimeras were removed with UCHIME (Edgar et al. 2011) and sequences were clustered into Molecular Operational Taxonomic Units (MOTUs) using swarm (Mahé et al. 2015), setting the threshold to d=13 (Siegenthaler et al. 2019, Atienza et al. 2020). This parameter sets the maximum distance, i.e. the maximum number of different nucleotides, which two random sequences may have in order to still be considered the same MOTU. Taxonomic assignment was performed with the BLASTN algorithm implemented in OBITOOLS against the MIDORI2 reference database (Leray et al. 2022) for COI sequences. In the same way, for 18S sequences, BLASTN was run against the PR^2^ database (Guillou, 2013). MIDORI2 and PR^2^ databases were selected, respectively, because of their specificity; the first contains quality-checked information on mitochondrial metazoan sequences, while the second is a combination of three interconnected 18S rRNA datasets, creating a Protist Ribosomal Reference database, useful for metabarcoding applications. Contaminants were removed using negative extraction controls, with the MICRODECON (version 1.0.2; McKnight et al. 2019), an R package specifically designed for identifying and removing contaminants. Final refinement of the dataset consisted of removing MOTUs with a <97% identity match with reference sequences. Non-target taxa (e.g., fungi, insecta, *Homo sapiens*, *Sus scrofa*, *Canis lupus*), as well as marine taxa occurring in the bait composition, that is *Salmo salar* and *Salvelinus* sp., were removed from the final dataset. During a preliminary inspection at the 18S dataset, taxa falling outside the intended target group (protists) were detected despite the use of protist-specific primers (Amaral-Zettler et al., 2009; Bradley et al., 2016). Therefore, metazoan taxa were removed from the final dataset, in accordance with previous metabarcoding studies (Leduc et al., 2019; Kumar et al., 2022). Additionally, some non-protist taxa (namely three Actinopteri) were likely due to misassignments, as 18S rRNA reference databases are mainly focused on protists or are less curated for metazoans (Di Capua et al., 2024).

### Data analysis

eDNA detections were considered as qualitative information, hence COI and 18S datasets were transformed from read abundances to presence/absence data using the *decostand* function from VEGAN R package (Oksanen et al., 2022).

#### Community composition

Community composition was analysed through non-metric multidimensional scaling (nMDS), via *metaMDS* function (VEGAN R package), applied using Jaccard distance, to visualize similarities among sites. A panel was plotted to show differences in sites distribution in a two-dimensional space. In the remaining panels of nMDS plot, for interpretative purposes, some taxa were aggregated into functional groups (discussed below and Table S1 and S2 in Supplementary materials) and divided into five trophic categories, according to their trophic level –primary producers (TL=1), primary consumers (TL=1.1–2.9), secondary consumers (TL=3–3.9), top-predators (TL≥4), detritivores and decomposers (taxa in the latter group were identified based on their diet composition) – to simplify visualization and highlight ecological patterns across samples.

The analysis of similarities (ANOSIM, *anosim* function from VEGAN R package) was conducted to test whether a significant difference existed between the communities identified in coastal and offshore samples. Additionally, an Indicator Species Analysis (IndVal, *multipatt* function, INDICSPECIES R package; De Cáceres et al., 2009) was conducted to evaluate whether certain species could be indicators of coastal or offshore communities.

#### Food web analysis

To evaluate the status of marine communities, a Molecular Ecological Network Analysis (MENA, Deng et al., 2012; Stat et al., 2019) approach was implemented. This process consists of three main steps: (i) reconstructing the alpha diversity of specific sites; (ii) mapping species interactions in terms of feeding relationships through literature review, and (iii) quantifying ecological network structure and properties. All the information obtained from literature review is available in the Supplementary materials (Table S1 and S2). The trophic level was first estimated using the function *ecology* from the FISHBASE R package, or, alternatively, when no information was returned, the *estimate* function from the same package. They collect information from widely used online databases: FishBase (Froese and Pauly, 2022) and SeaLifeBase (Palomares, 2022). When no information was available in existing databases, we developed a custom function in R using the RGLOBI package (Poelen et al., 2023), which allowed us to estimate the trophic level of target taxa based on their reported diet. The RGLOBI package provides a list of species that are “eaten by” or “preyed on” the focal taxon. The trophic level of each taxon was defined as one plus the mean trophic level of the taxa comprising its diet. For basal species, our literature review allowed us to distinguish between autotrophs (whose trophic level is equal to one, as they are primary producers) and heterotrophs or mixotrophs, for whom the above method was applied. Some taxa were aggregated into functional groups, defined as groups of species that have the same role in the trophic web (Table S1 and S2 in Supplementary materials). The trophic level of each group was calculated as the mean of the trophic levels of its constituent species. Subsequently, local food webs were reconstructed using the CHEDDAR R package (Hudson et al., 2013); a food web for each site and one for each group of samples (coastal and offshore) was plotted. The food webs were visually explored and then characterized using the following network indicators (Table 3): (1) Species Richness; (2) Number of links; (3) Direct connectance; and (4) Generality. These indicators were used to summarise the food web structure (topological characteristics) and network complexity for each local food web.

**Table 3.** Full names and codes of the food web indicators used in this study, with their explanation and their ecological meanings.

| Food web indicator | Explanation | Interpretation and References |
| --- | --- | --- |
| Species Richness (N) | Number of taxa (nodes) in a food web | Diversity degree in the community. (May, 1974; Tilman, 1996) |
| Number of links (L) | Number of trophic links in the community | Number of energy transfers into the community. (Dunne et al., 2002) |
| Direct<br>connectance<br>(C) | Number of trophic links/<br>number of $n$ nodes<sup>2</sup>,<br>including cannibalistic<br>links. Measure the proportion of realised<br>interactions<br>among all the possible<br>ones in a network | Lower connectance value can<br>reveal a decrease in food web<br>robustness. (Martinez, 1992;<br>Dunne et al., 2002) |
| Generality<br>(G) | Average number of prey<br>per predator<br>(Schoener, 1989) | Indicates if the system<br>contains<br>more generalist or<br>specialist species. |

## RESULTS

### Sequencing output

After bioinformatic processing, a total of 103,259 reads were obtained from COI sequencing and 378,818 reads from 18S. Metazoans were removed from the 18S dataset because the reference databases for this barcode region focus primarily on protists, which could result in an erroneous classification of metazoan taxa.

Following decontamination, data filtering and non-target taxa removal, the final dataset comprised 111 taxa belonging to 21 phyla for COI, and 95 taxa distributed across 13 phyla for 18S. Taxonomic assignment varied in resolution: for COI, 71 taxa were identified at the species level, 34 at genus, 3 at family, 2 at order and 1 at class level; while for 18S, 27 taxa were resolved at the species level, 47 at genus, 8 at family, 9 at order and 4 at class level (Figure 2).

**Figure 2.**
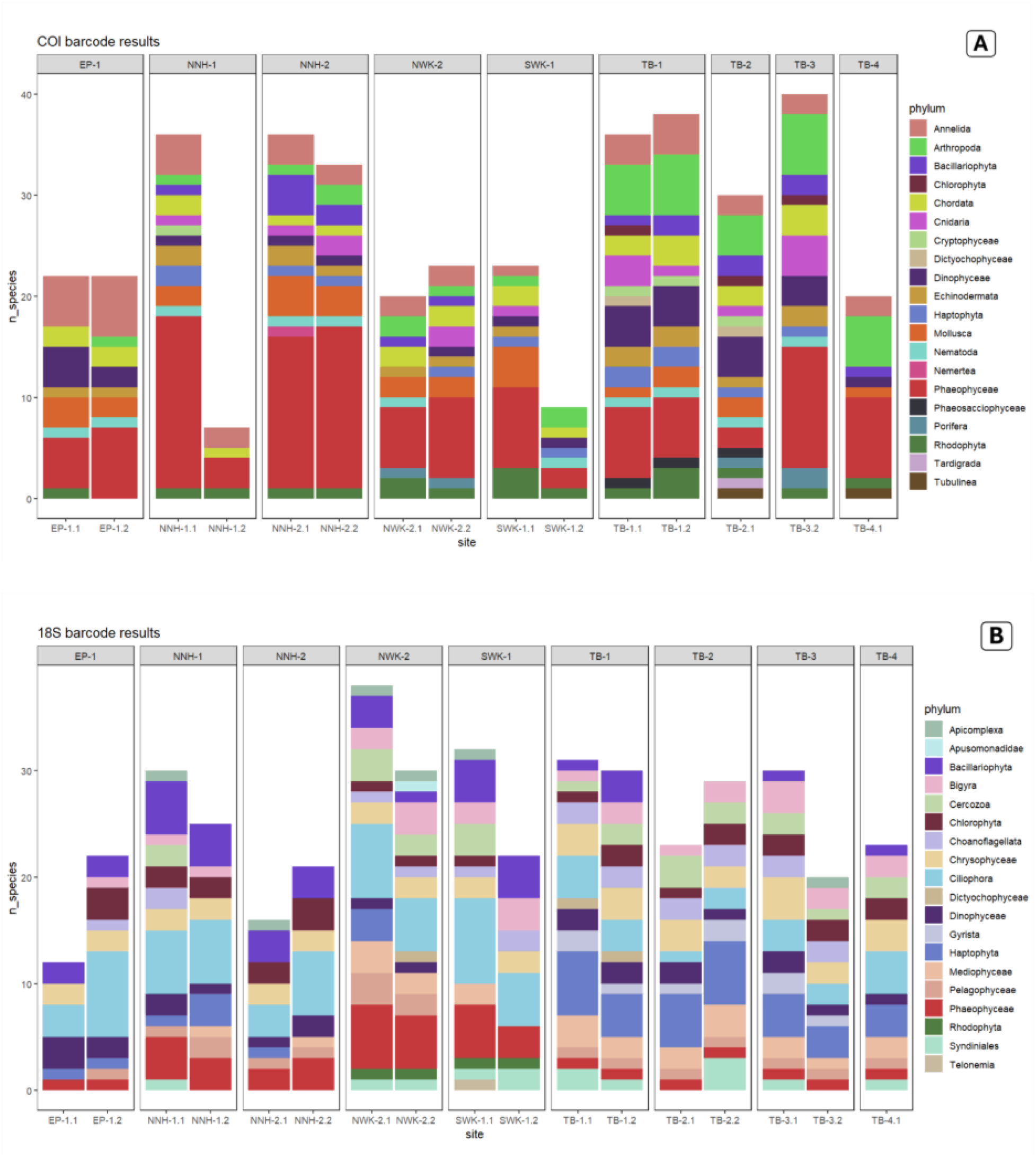
For each sample reported on the x-axis, the number of taxa (y-axis) is shown grouped by phylum, for the COI (A) and 18S (B) barcode regions.

In the COI dataset, the Phaeophyceae phylum showed 28 amplified taxa, including microalgae (like Haptophyta and Dictyochophyceae), macroalgae (as Phaeophyceae, Phaeosacciophyceae and Rhodophyta) and diatoms from the Bacillariophyceae. Other microorganisms belonged to: Dinophyceae (dinoflagellates), Tardigrada, and Tubulinea (amoeba).

Chordata were represented by three Arctic fish species: *Boreogadus saida*, a key forage fish in Arctic ecosystems; *Myoxocephalus scorpius*, a benthic sculpin; and *Liparis liparis*, a snailfish commonly associated with cold-water environments. In addition, marine mammals were identified, including *Phoca groenlandica* (harp seal), *Delphinapterus leucas* (beluga whale), and *Balaenoptera acutorostrata* (minke whale), all species characteristic of Arctic marine habitats. Finally, one Ascidiacea, *Halocynthia pyriformis*, was also detected.

Other phyla were also well represented: Annelida (13 taxa), spanning both Polychaeta and Clitellata and covering a variety of marine worm families, Arthropoda (10 taxa), mostly crustaceans, including Malacostraca, Thecostraca, and Hexanauplia, and Mollusca, with 4 taxa each from Bivalvia and Gastropoda. Echinoderms were also identified, including four species across Echinoidea (sea urchins), Holothuroidea (sea cucumbers), and Ophiuroidea (brittle stars). Additionally, 8 taxa belonging to four classes of Cnidaria (i.e., Anthozoa, Hydrozoa, Scyphozoa, and Staurozoa) were retrieved, reflecting a broad diversity of cnidarian lineages captured in the dataset.

Lastly, Nematoda and Nemertea were also detected, although represented by a limited number of taxa. These included the nematode genus *Pseudochromadora* and the ribbon worm *Malacobdella grossa*.

As expected, from the 18S dataset numerous microorganisms were retrieved. These included a wide array of algal taxa, such as Cryptophyceae, Chrysophyceae, Pelagophyceae, Mediophyceae, Dictyochophyceae, and Haptophyta, as well as diatoms from the phylum Bacillariophyta. Also macroalgal species belonging to Chlorophyta, Rhodophyta and Phaeophyceae were registered. A conspicuous representation of Alveolata was found, dominated by Ciliophora, with 20 taxa, in addition to seven taxa from Dinophyceae and Syndiniales, the latter consisting of parasitic dinoflagellates infecting crustaceans, fishes, cnidarians, and other protists. Finally, the dataset also revealed microscopic eukaryotes, such as Cercozoa and Bigyra.

### Community composition

The non-metric multidimensional scaling (nMDS) revealed a clear separation between coastal and offshore samples, which clustered in distinct regions of the ordination (Figure 3A). This separation was primarily driven by a gradient along the first axis (NMDS1). Additionally, the coastal samples collected at the fjord entrance (i.e., NWK) clustered closer to the TB samples.

**Figure 3.**
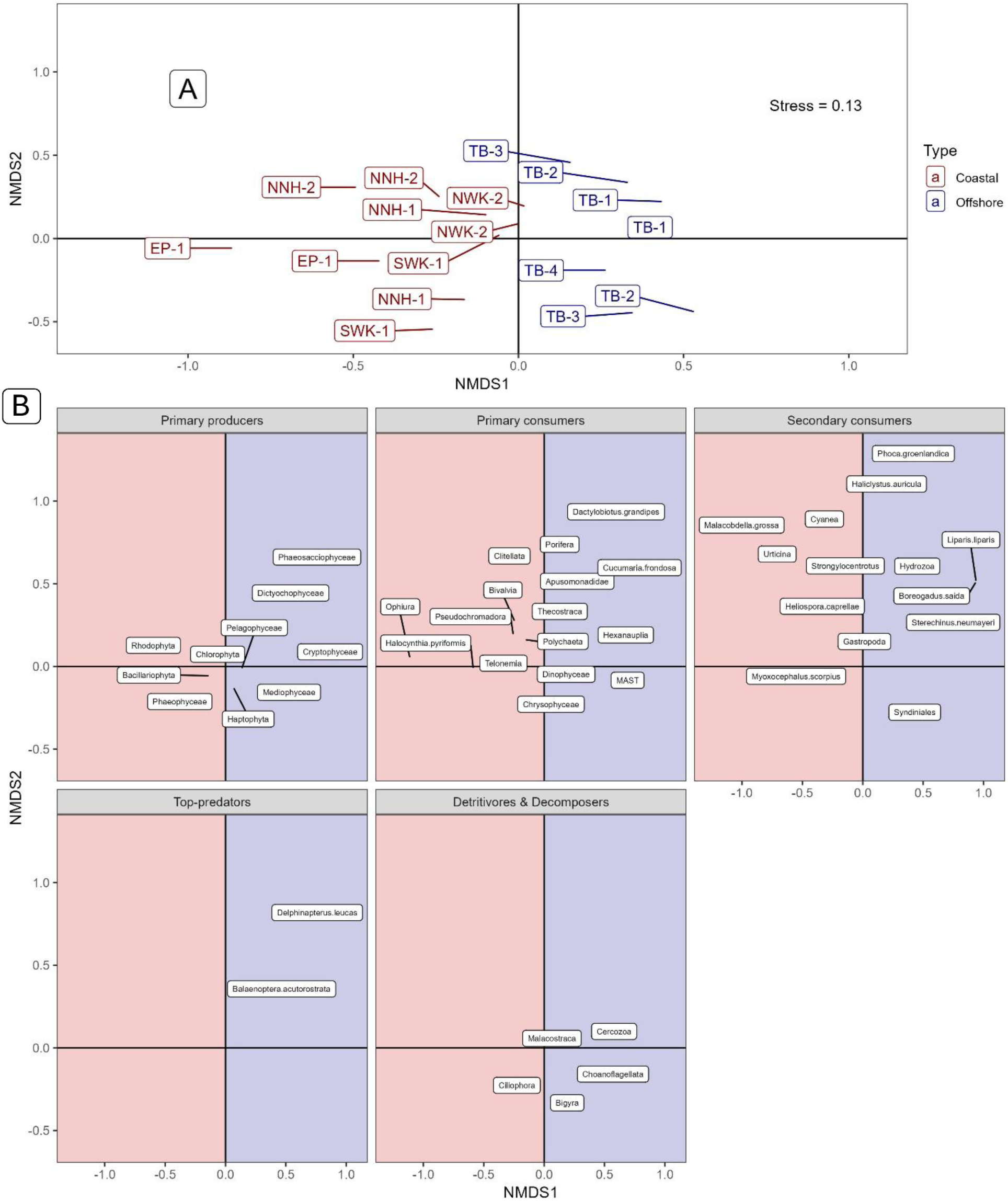
(A) Non-metric multidimensional scaling (nMDS) plot showing both laboratory replicates, for each sampling site. (B) nMDS plots based on aggregated taxa, grouped into five trophic categories.

In panels B, species are plotted according to their distribution across the nMDS ordination. First, primary producers such as microalgae and diatoms (phytoplankton) clustered predominantly in the offshore samples, whereas macroalgal taxa belonging to Rhodophyta and Phaeophyceae are mainly located in the coastal area. Concerning the primary consumers, zooplanktonic taxa, like *Calanus* species (Hexanauplia), are positioned in the offshore domain as they feed on phytoplankton. Conversely, benthic taxa such as Clitellata and Amphipoda (Malacostraca) are concentrated in the coastal area, as they exploit phytodetrital and macroalgal-based carbon sources (e.g. Rhodophyta and Phaeophyceae). The secondary consumers showed a similar pattern. For instance, the benthic-associated fish *Myoxocephalus scorpius* clusters in the same domain like its main prey items, such as *Bivalvia* and partially *Gastropoda*, that is in the coastal region of the plot. Similarly, *Phoca groenlandica*, located in the offshore samples, clusters together with its prey (*B. saida* and *L. liparis*). Top predators exhibit a broader diet, including both fishes and invertebrates (*Polychaeta*, *Malacostraca*, *Thecostraca*, *Porifera*, and mollusks), which are distributed across both domains. As for detritivores and decomposers, they are found in both environments but the higher species richness of Ciliophora gives this group greater weight in shaping the observed distribution.

The ANOSIM test confirmed a positive and significant difference between the two sampling groups (R-values=0.7, *p*=0.001; Table S3 in Supplementary materials). Furthermore, the dissimilarity rank between groups (91.25%) was higher than those observed within groups (51.50% for coastal and 22% for offshore units), supporting the robustness of this distinction.

The association degree between a species and a sampling group was determined by IndVal (Table S4 in Supplementary materials). The coastal group was characterized by 16 associated species, including high-indicator taxa such as the diatom *Cylindrotheca closterium* (stat=0.998, p-value=0.001) and the Rhodophyta group (stat=0.941, p-value=0.005). Additionally, *Myoxocephalus scorpius* was found to be associated with the coastal group. In contrast, the offshore group showed a much higher number of associated species (31 total), dominated by key open-water organisms. The copepod *Calanus* emerged as an indicator species for this group (stat=1.000, p-value=0.001), alongside several microalgae such as *Micromonas polaris*, *Chrysochromulina* and the marine stramenophiles *MAST-3I* sp. As well as other algal groups like Chlorophyta, Haptophyta and Mediophyceae, all of them showed an association value close to one, making them well-fitted indicators for the offshore group.

### Food web analysis

Among all taxa and functional groups considered, a total of 51 nodes were included in the reconstructed food web (Table S5 in Supplementary materials). In addition to the amplified taxa, basal resources such as benthic detritus, particulate organic matter (POM), bacteria, and inorganic components were also incorporated, following Cicala et al. (2024).

Primary producers were represented by several algal groups, including Bacillariophyta, Chlorophyta, Cryptophyceae, Dictyochophyceae, Haptophyta, Mediophyceae, Pelagophyceae, Phaeophyceae, Phaeosacciophyceae, and Rhodophyta. These autotrophic and photosynthetic taxa, ranging from micro- to macro-scopic forms, constitute part of the self-feeding community that underpins marine food webs.

Primary consumers (trophic level 2–2.9) were represented by different functional groups. These included protists such as Telonemia, Apusomonada, Bigyra and marine stramenopiles (MAST), as well as metazoans like Clitellata, Porifera, Thecostraca, Polychaeta, Malacostraca, Hexanauplia, Bivalvia, and Ascidiacea. Heterotrophic dinoflagellates (Dinophyceae) and mixotrophic Chrysophyceae were also placed in this category, reflecting the metabolic plasticity of several consumer groups.

Secondary consumers included taxa whose ecological roles are often more complex than their phylogenetic affiliations might suggest. For example, *Heliospora caprellae* and members of Syndiniales, although belonging to groups typically associated with lower trophic levels, were identified as parasites of marine crustaceans and thus occupy higher positions in the food web. Other secondary consumers included fish, such as *Boreogadus saida*, benthic or pelagic invertebrates, like Gastropoda, Hydrozoa, and *Malacobdella grossa* (a commensal nemertean worm), as well as cnidarians such as *Haliclystus auricula*. Echinoderms like brittle stars (*Ophiura sp.*) and sea urchins (*Sterechinus neumayeri*, *Strongylocentrotus sp.*), were also part of this group.

Apical species included the harp seal *Phoca groenlandica,* the northern minke whale *Balaenoptera acutorostrata* and the beluga whale, *Delphinapterus leucas,* with respectively assigned trophic levels of 3.69, 4.2 and 4.25. These species were found only in offshore sites. Among the coastal sites, the main predators were represented by benthic and demersal fish, such as the snailfish (*Liparis liparis*) and the sculpin (*Myoxocephalus scorpius*), with trophic levels of 3.59 and 3.69, as well as large cnidarians like *Urticina* sp. and the jellyfish genus *Cyanea* sp., which reached trophic levels of 3.64 and 3.62.

Local food webs showed marked variability in their structural indicators (Table 4 and Figure 4). Sites NWK-2 and TB-1 displayed the highest number of nodes (34 and 37, respectively), whereas the lowest richness was recorded at site EP-1, with only 21 nodes. Accordingly, the number of links ranged from 48 (EP-1) to 115 (TB-1), spanning across four trophic levels. As expected, sites with the greatest number of nodes also tended to exhibit the highest number of links (e.g., TB-1, NWK-2).

**Figure 4.**
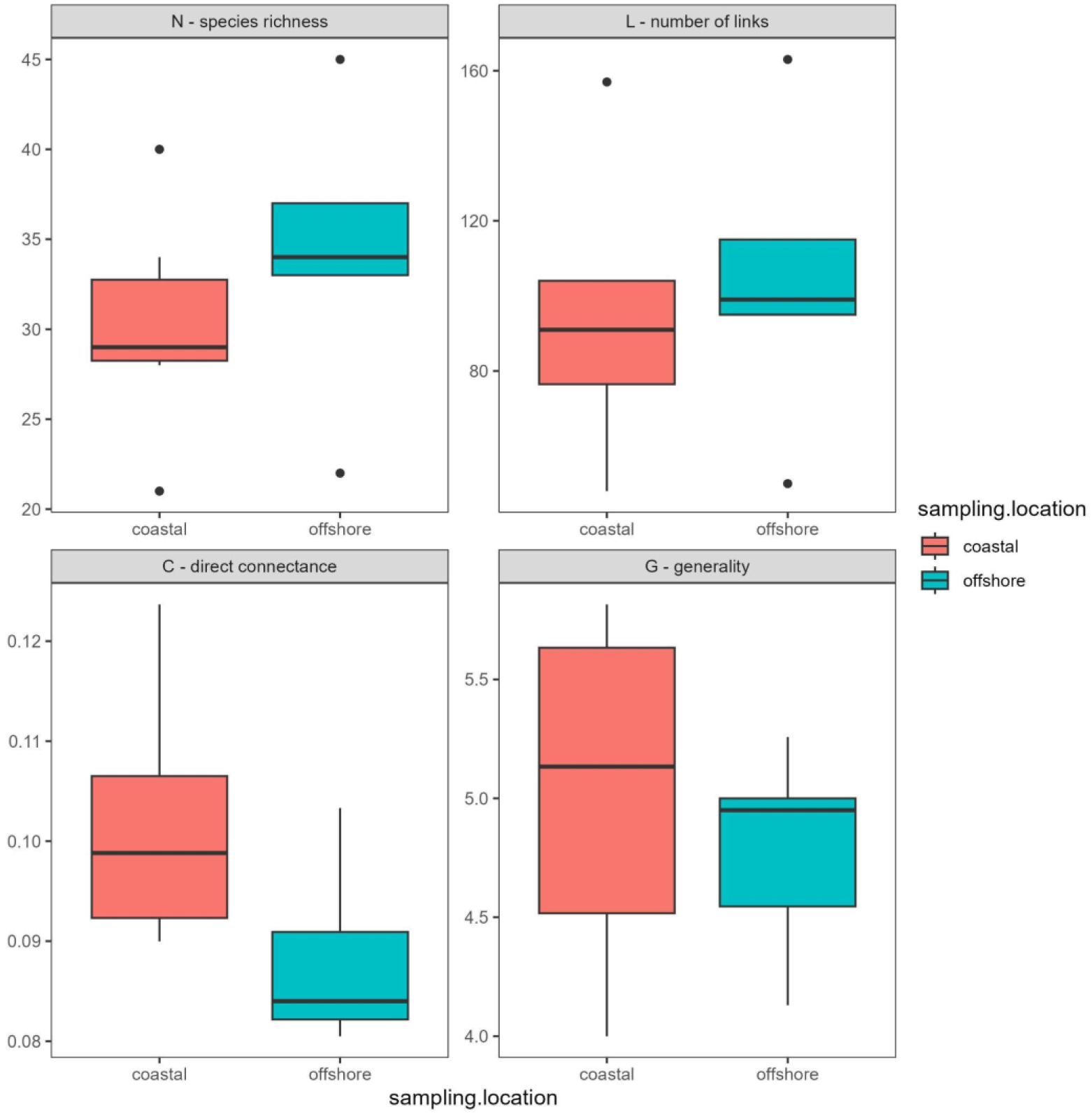
Overview of food web indicators for coastal and offshore sites.

**Table 4.**
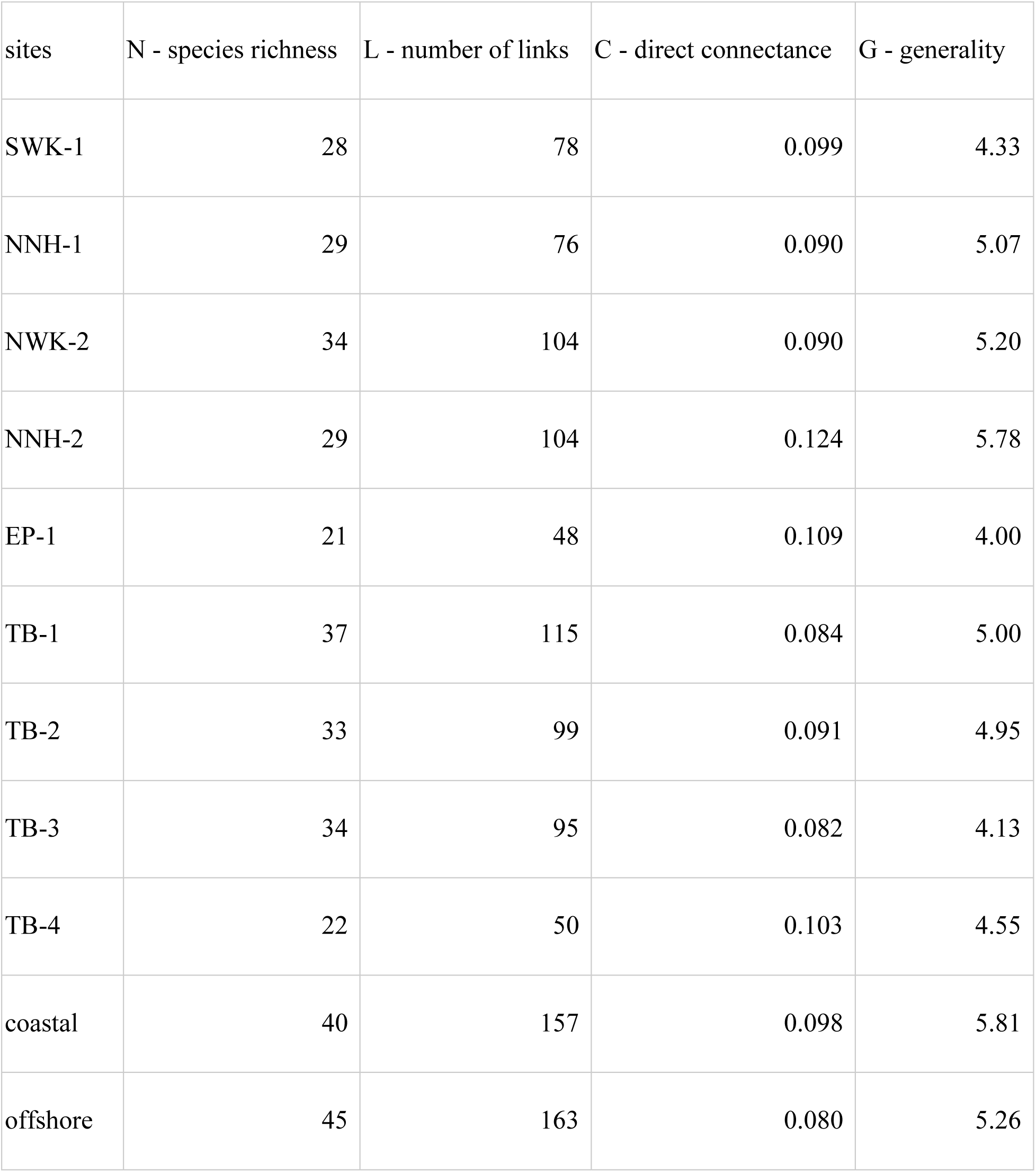
For each site, the four trophic indicators calculated are reported.

Connectance values, however, did not follow the same pattern. Site TB-3, which despite hosting a considerable number of nodes (N = 34) recorded the lowest connectance value (C = 0.082). Conversely, site EP-1 and TB-4, even though they had the lowest number of nodes and links, presented higher connectance values (around 0.1 for both sites). Overall, the highest connectance value (0.124) was that of site NNH-2, which also showed the highest generality value (5.76). The lowest generality value was found in EP-1 site (G = 4.00), this location indeed maintained a low number of nodes and links. TB-4, while sharing the same patterns of N and L as EP-1, displayed a wider dietary range for its species, with a G value of 4.55, moreover highlighting a more tightly connected network (C = 0.103) despite its limited species richness. The average number of prey per predator across the complete food web was 5.67, with local values ranging from 4.00 in EP-1 to 5.78 in NNH-2.

When aggregating the four offshore and five coastal sites, offshore sites overall showed higher species richness compared to coastal sites. The most evident differences emerged at the top of the food webs. Apical predators such as the minke whale *Balaenoptera acutorostrata*, the beluga whale *Delphinapterus leucas*, and the harp seal *Phoca groenlandica* were detected exclusively in offshore sites (Figure 5). In contrast, the apex of the coastal sites’ food web was represented by fewer predators, namely the fish *Myoxocephalus scorpius* and the cnidarians *Cyanea sp.* and *Urticina sp.* (Figure 6). At lower trophic levels, species composition was broadly similar between the two groups, with the exception of the holothurian *Cucumaria frondosa*, absent from coastal sites, and the ascidian *Halocynthia pyriformis* missing from the offshore domain.

**Figure 5.**
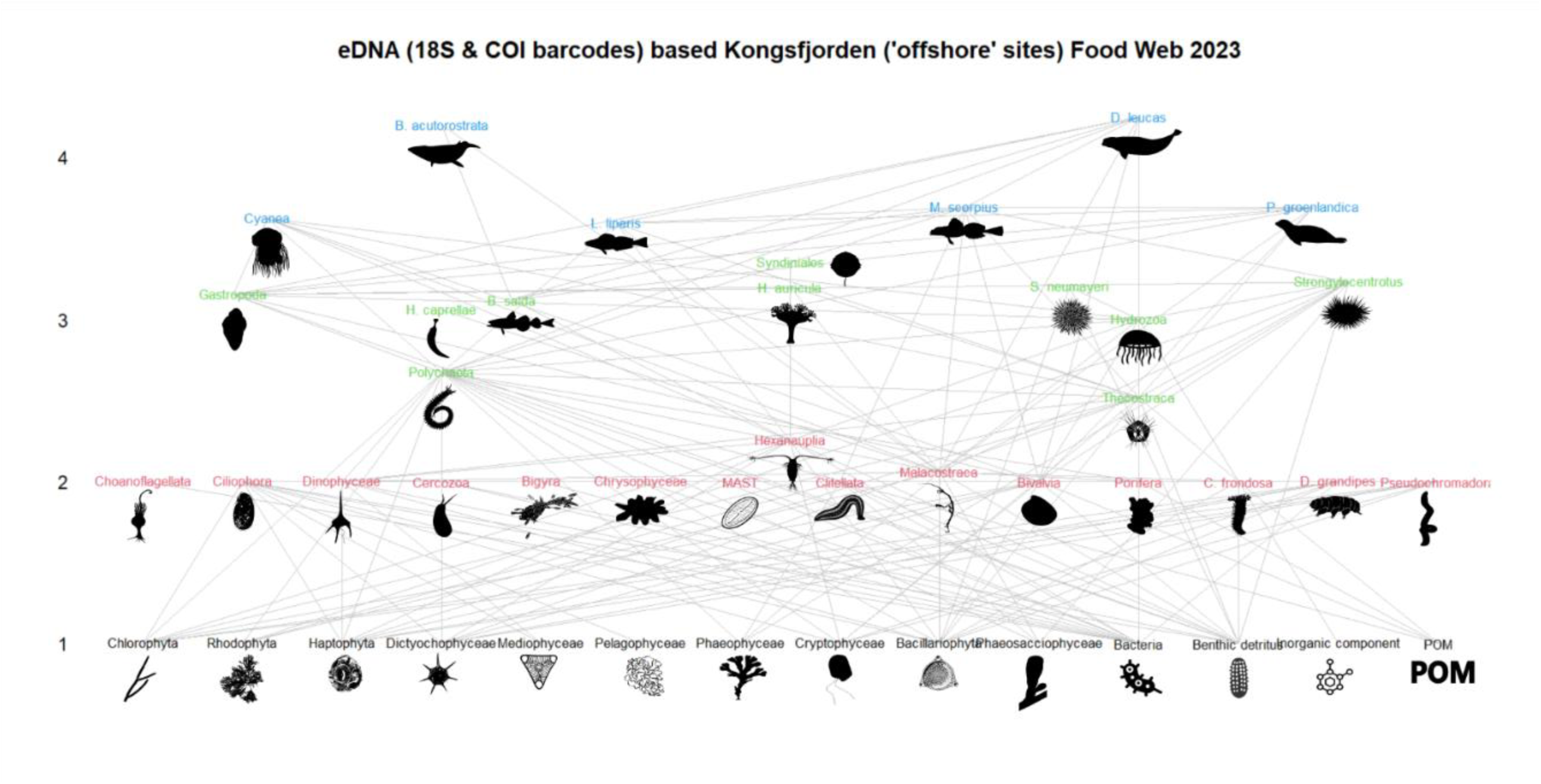
Food web from the aggregated offshore and coastal sites (see also Figure S10 in Supplementary Materials). Values on the y-axis correspond to trophic levels. Only the taxa present in the location are plotted. Site-specific food webs can be found in Supplementary Materials (Figures S1 to S9).

**Figure 6.**
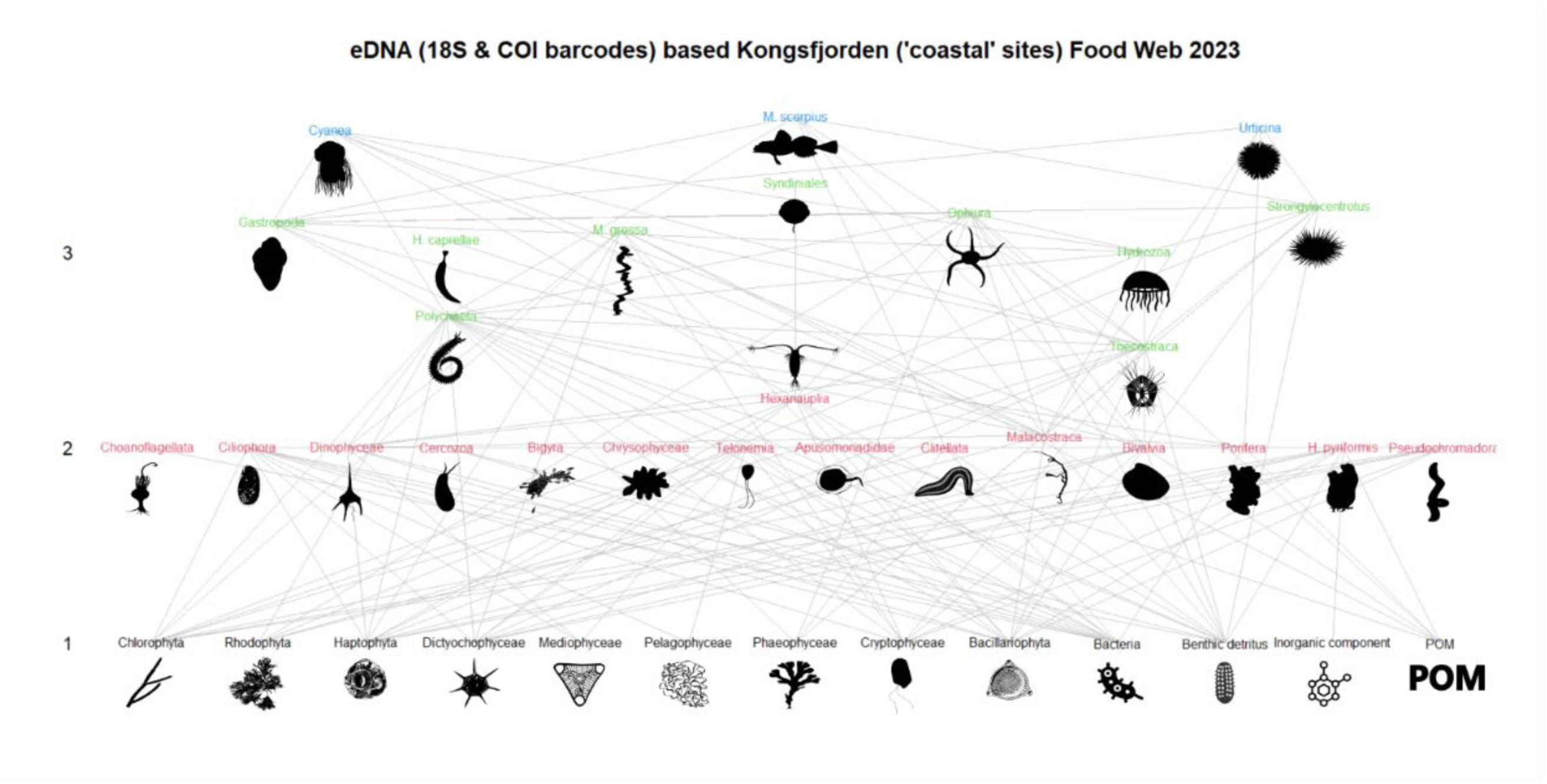
Food web from the aggregated offshore and coastal sites (see also Figure S10 in Supplementary Materials). Values on the y-axis correspond to trophic levels. Only the taxa present in the location are plotted. Site-specific food webs can be found in Supplementary Materials (Figures S1 to S9).

Generality values appeared to be higher in coastal sites compared to offshore ones (5.77 vs 5.20), in line with the higher connectance observed locally (0.099 vs 0.081). This suggests that predators in coastal sites tend to exploit a wider range of prey items. For instance, within our dataset *Balaenoptera acutorostrata* had only two links (*Boreogadus saida* and Euphausiacea, included in Malacostraca), while *M. scorpius* displayed a more generalist diet including, between other items, *B. saida*, *L. liparis*, and *Strongylocentrotus* sp.

## DISCUSSION

### Sequencing output

As the Arctic area undergoes unprecedented shifts, understanding the structural integrity of its marine life is more critical than ever. Thus, the present study aimed at exploring the Kongsfjorden ecosystem through the application of eDNA metabarcoding approach, that is transforming genetic signals into a detailed map of the fjord’s food web and ecological equilibria. Each of the used gene markers displayed differences in taxonomic resolution. COI provided a finer resolution, most frequently allowing taxonomic assignment to the species level, while 18S gene showed a coarser resolution, with most taxa classified at genus or higher ranks. Identification to the species level may be lacking due to the absence of the respective sequences in reference databases and/or because maybe 18S and COI marker regions showing insufficient polymorphisms among species within a genus to properly distinguish them (van den Heuvel-Greve et al., 2021).

Our aim was to comprehensively identify community composition rather than compare the diversity and taxonomy of different molecular markers. The choice of using the 18S barcode was made in order to explore the unicellular eukaryotes that inhabit the study area. Hence, two primers targeting V8-V9 regions of the gene were chosen, as they were designed to detect the aquatic microbial community (Amaral-Zettler et al., 2009; Bradley et al., 2016).

Despite being a marker targeting metazoans, COI also retrieved a considerable fraction of algal taxa, as it is known that this primer shows non-specific amplification as well (Collins et al., 2019). This could be linked to the presence of great biomasses of algal cells and cellular fragments in the water column, especially in productive shallow Arctic coastal waters, where phytoplankton can dominate the DNA pool and thus be co-amplified by universal primers (Specchia et al., 2022).

Taken together, these results underline how the combined use of multiple markers provides complementary insights. Indeed, Ficetola and Taberlet (2023) have proposed emerging approaches that would allow more holistic reconstruction of ecological communities, like the combination of universal and specific primer pairs or the use of shotgun sequencing. Concerning our study, COI enhanced taxonomic resolution for animals, while 18S captured a wider spectrum of microbial eukaryotes, though with lower resolution. Notably, both barcodes also detected parasitic or symbiotic organisms (e.g., Nemertea’s *Malacobdella grossa*, some Clitellata worms, Syndinium, *Heliospora caprellae*), highlighting the ability of eDNA to reveal ecologically important taxa that are often overlooked by traditional survey methods.

However, eDNA metabarcoding still has to face some limitations, regarding the availability of sufficient and unambiguous DNA sequences in reference databases, as well as the development of markers that allow a better coverage of all key taxonomic groups within the wide marine diversity (Lacoursière-Roussel et al., 2018; Van der Loos and Nijland 2020). This was evident when first investigating our datasets and it is even more accentuated for those under-studied organisms such as those inhabiting remote regions like the Arctic (Leduc et al., 2019).

Regarding sample quality, a subset of samples (one replica each from TB-2, TB-3 and TB-4 samples in the COI dataset, and one replica from TB-4 sample in the 18S dataset) had to be discarded due to insufficient DNA content. Several factors could have contributed to this outcome. One possible explanation could be the immersion between sampling types: offshore samples were deployed for about one hour, whereas coastal samples remained in the water for 24 hours. Such variation could have affected DNA yields, reflecting differences linked to sampling duration and strategy, as these two factors are known to influence successful DNA recovery (Jackman et al., 2024). However, the extent to which sampling methods with *metaprobes* affect DNA recovery still requires further investigation. Moreover, these outcomes emphasize how sampling design and sample preservation can strongly influence DNA recovery and the overall reliability of downstream analyses (Yamanaka et al., 2021).

### Ecology and community composition

The nMDS analysis revealed a clear separation between coastal and offshore samples, reflecting the distinct environmental characteristics of the two areas. Coastal sites, located in the harbour area of Ny-Ålesund, are shallower and more influenced by anthropogenic activities, while offshore samples were collected in the central part of Kongsfjorden, characterized by deeper waters and reduced human impact. NWK samples clustered closer to TB, reflecting the environmental similarity of the two habitats, both representing more open areas within the fjord.

This separation was also influenced by the sampling method: coastal samples, collected with set traps, captured a more benthic community, whereas offshore samples obtained through towed sampling were richer in pelagic taxa. The distribution of functional groups supports this pattern: primary producers (micro- and macroalgae), primary and secondary consumers, and detritivores and decomposers are represented across both domains, yet a higher species richness of an aggregated taxa gives that group greater weight in shaping the observed distribution. The detection of pelagic top predators (*Balaenoptera acutorostrata*, *Delphinapterus leucas*) exclusively in offshore sites aligns with their lower tolerance to high anthropogenic disturbance, which typically characterizes more coastal environments, in particular as it can mask up cetacean communication signals (Lechwar et al., 2023) and shows reliability and precision of eDNA in depicting spatial patterns.

The relationships between consumers and resources identified by the literature review appear to follow a consistent pattern, namely that a consumer and its resources are located in the same area of the nMDS. The distribution of resources consumed by top predators also reflects the coupling of distinct trophic channels described by McMeans et al. (2013). Phytoplanktonic taxa (microalgae and diatoms) dominated the offshore domain, whereas macroalgae (Phaeophyceae, Rhodophyta) characterized the coastal domain. This spatial organization mirrors the distinction between the grazing and detrital food chains: offshore, the grazing pathway predominates, driven by phytoplankton–zooplankton–fish–mammal interactions; coastal systems, instead, are dominated by benthic detrital pathways involving decomposers and detritivores such as Ciliophora and Malacostraca (namely the two Amphipoda taxa that make up this group), and the apex of the web is dominated by benthic-associated *M. scorpius*. Even though at lower trophic levels this compartmentalization remains, upper trophic levels integrate resources from both coastal and offshore food chains, acting as couplers of distinct trophic pathways which grant stability to ecosystems as reported by Rooney et al. (2006) and McMeans et al. (2013).

Results from ANOSIM and IndVal analyses confirmed the presence of distinct assemblages characterizing coastal and offshore environments. The identification of indicator taxa, which were completely distinct and exclusive for the two groups, demonstrates the sensitivity of eDNA metabarcoding to detect community-level differences even at small spatial scales.

It is important to state that the food web reconstruction presented here is qualitative, as it is based on presence/absence data without biomass or energy flow estimates. Nevertheless, the resulting network effectively captured the main trophic structure and ecological relationships. Some biases remain, as trophic links inferred from literature may oversimplify real feeding interactions (D’Alessandro and Mariani, 2021); however, the approach provides an integrative view of community organization without the invasiveness or effort of traditional methods.

Regarding trophic indicators, the pattern observed in species richness (N) and trophic links (L) is consistent with previous findings showing that these network indicators, particularly species richness, reflect different levels of anthropogenic impact (D’Alessandro & Mariani, 2021). In our case, both N and L were lower in coastal sites, those located near the harbour, where human influence is likely stronger, whereas higher values were recorded offshore. This mirrors the difference in community composition, which likewise differed between coastal and offshore domains, suggesting that anthropogenic disturbance may reduce the number of trophic groups and the density of interactions within the food web (D’Alessandro and Mariani, 2021).

Connectance (C), in turn, is known to decrease under stressful conditions (Safi et al., 2019). Offshore communities showed lower C values together with lower generality values, indicating narrower dietary ranges and a greater degree of trophic specialization. Such specialization likely intensifies competition for a limited set of resources, resulting in a less stable network in the offshore community (Le Guen et al., 2025). In contrast, coastal communities displayed higher C values, consistent with their more generalist feeding strategies. This broader resource use may alleviate competitive pressure and allow a greater proportion of potential trophic links to be realised, despite the higher anthropogenic impact in these areas (D’Alessandro and Mariani, 2021). In general, it is important to study connectance values to determine what provides stability to an ecosystem and what, on the other hand, causes stress. Moreover, describing trophic relationships can help to predict the relative stability of the system when confronted with species introductions/extinctions, altered productivity patterns and other natural and human-induced system changes (Renaud et al., 2011).

eDNA metabarcoding proved to be a powerful complementary tool for food web reconstruction, offering high taxonomic resolution, reproducibility, and spatial coverage. Its integration with traditional approaches could provide a comprehensive framework for monitoring ecosystem structure and stability in remote and rapidly changing environments like the Arctic.

## CONCLUSION

This study demonstrates that eDNA metabarcoding, and its application with *metaprobes*, enabled us to successfully reconstruct trophic webs in every site and to detect differences in community composition when considering more domains of an ecosystem. This allowed us to obtain information on local biodiversity by boosting data acquisition and revealing community composition at high resolution.

In a rapidly warming Arctic undergoing profound ecological shifts, this approach represents a valid tool to investigate ecosystem stability and, specifically, variations in community composition that could alter trophic webs’ structure. Its rapid application, moreover, can provide the baseline for long-term monitoring studies, allowing comparison of trophic networks over time. Despite the qualitative nature of the presented trophic webs, they can still provide a realistic portrayal of marine communities through significantly reduced costs. Integration with traditional methods (e.g., stomach content analysis and stable isotope analysis) could allow for a more accurate representation of networks as they consider energy flows between trophic compartments. Overall, eDNA-based food web analysis offers a powerful and practical framework for early detection of ecosystem changes driven by climate warming and anthropogenic stressors, supporting informed management and conservation strategies in Arctic marine ecosystems.

## Authors’ Contributions

CP: Conceptualization, Data curation, Formal analysis, Visualization, Writing – original draft. NM and SG: Formal analysis, Laboratory analysis, Data interpretation, Writing – review & editing.

TR: Methodology, Data interpretation, Supervision, Writing – review & editing.

MA and FF: Investigation, Sampling, Resources.

GDM, ADA and AG: Labooratory analysis, Data interpretation, Writing – review & editing,

AP: Conceptualization, Project administration, Funding acquisition, Supervision, Writing – review & editing.

## Data Archiving Statement

The raw sequence data for the DNA metabarcoding experiment are deposited in the NCBI Sequence Read Archive (Bioproject accession PRJNA1391418).

## Acknowledgements

The authors acknowledge the Arctic research base Dirigibile Italia (Ny-Ålesund, Svalbard), managed by CNR-ISP, for the logistical and operational support provided during the VECNA 2023 field activities. Special thanks are extended to the station leader, Simonetta Montaguti, for her coordination and assistance during the survey. This research was financially supported by the Institute for Biological Resources and Marine Biotechnology (CNR-IRBIM), which contributed to the implementation of the VECNA 2023 sampling survey.

